# The Structural Basis of IKs Ion-Channel Activation: Mechanistic Insights from Molecular Simulations

**DOI:** 10.1101/275180

**Authors:** Smiruthi Ramasubramanian, Yoram Rudy

## Abstract

Relating ion-channel (iCh) structural dynamics to physiological function remains a challenge. Current experimental and computational techniques have limited ability to explore this relationship in atomistic detail over physiological timescales. A framework associating iCh structure to function is necessary for elucidating normal and disease mechanisms. We formulated a modeling schema that overcomes the limitations of current methods through applications of Artificial Intelligence Machine Learning (ML). Using this approach, we studied molecular processes that underlie human IKs voltage mediated gating. IKs malfunction underlies many debilitating and life-threatening diseases. Molecular components of IKs that underlie its electrophysiological function include KCNQ1 (pore forming tetramer) and KCNE1 (auxiliary subunit). Simulations, using the IKs structure-function model, reproduced experimentally recorded saturation of gating charge displacement at positive membrane voltages, two-step voltage sensor (VS) movement shown by fluorescence, iCh gating statistics, and current-voltage (I-V) relationship. New mechanistic insights include - (1) pore energy profile determines iCh subconductance (SC), (2) entire protein structure, not limited to the pore, contributes to pore energy and channel SC, (3) interactions with KCNE1 result in two distinct VS movements, causing gating charge saturation at positive membrane voltages and current activation delay, and (4) flexible coupling between VS and pore permits pore opening at lower VS positions, resulting in sequential gating. The new modeling approach is applicable to atomistic scale studies of other proteins on timescales of physiological function.

## INTRODUCTION

A voltage-gated ion-channel (iCh), in response to membrane voltage (*V_m_*), undergoes atomistic structural changes that are stochastic in nature and underlie its physiological function as a transmembrane (TM) charge carrier. Recent advances in experimental techniques (1) provide considerable structural details that can be used to simulate atomistic structural dynamics. Relating these dynamics to the physiological function of proteins, including iCh, remains a major objective and difficult challenge in the field of biophysics. In principle, computational techniques can be used to simulate structure-based iCh function in order to explain its mechanism (2, 3). However, even with customized hardware, simulations of 1 msec gating dynamics take months and require non-physiological conditions, e.g. *V_m_* of 500mV (4, 5). Therefore, direct computation of iCh conformational changes during gating, with atomistic structural details and over a sufficiently long timescale (tens of msecs) is impractical.

In this paper, we introduce a novel methodology using Machine Learning (ML), for modelling iCh with structural and functional details. The method can be used to probe mechanisms that govern ionic conductance and voltage mediated gating at sufficient atomistic resolution over physiological timescales. The protein archetype we chose is the slow delayed rectifier IKs, a human voltage-gated potassium iCh. IKs is composed of a KCNQ1 pore forming tetramer modulated by KCNE1 segments. It plays an important role in cardiac action potential (AP) repolarization during *β*-adrenergic stimulation and participates in cardiac AP rate dependent adaptation (6, 7). IKs mutations are implicated in the most common congenital long QT syndrome Type 1 and in severe neurological disorders such as deafness and epilepsy (8). Additionally, this channel poses many structure-function molecular properties of interest, such as multiple subconductance (SC) levels, two-distinct gating conformational movements, voltage sensor (VS) or S4 movement that precedes ionic current, gating charge (GC) saturation at positive *V_m_*, and sequential gating (9–11). The new ML-based modelling approach simulates experimental data with high accuracy and reproduces these properties; it provides new mechanistic insights on the molecular basis of iCh activation under physiological conditions.

## METHODS

A short overview of simulating structural dynamics and electrophysiological function is provided below; details are included in Supplemental Information (SI) Section 1 and 2. The energy landscape was constructed to simulate IKs structural changes in response to *V_m_*. If done explicitly, this process requires computing energy of all possible IKs conformations. This is an impossible task, given the vast number of degrees of freedom. Previous modeling overcame these difficulties by retaining only large backbone movement with limited degrees of freedom, assuming tetrameric symmetry and reducing simulations to KCNQ1 TM segments (12–15). Here, we apply ML to simulate physiological IKs behavior at the atomistic scale, without the approximations and simplifications of previous work.

### Structure to Function

The initial IKs structure was constructed from experimental data using computational modeling (Figure 1); the specifics are detailed in SI Section 1. Extensive *de novo* sampling of IKs conformational space (library; ~3,000,000 conformations) was performed without any dimension reduction, applying physiological and experimental constraints. MODELLER (16) was used to minimize energy and remove steric clashes. Selected IKs attributes (features) that alter protein energy were extracted from the library, along with the computed energy of each corresponding structure (17, 18). Using these data, the ML algorithm (19) was trained to predict IKs energy of structures outside the library. With this approach, the energy landscape covered the entire IKs conformational space associated with activation. If desired, the structure for any point on the energy landscape could be retrieved. CHARMM force-fields were used for energy calculations (20).

**Figure 1:**
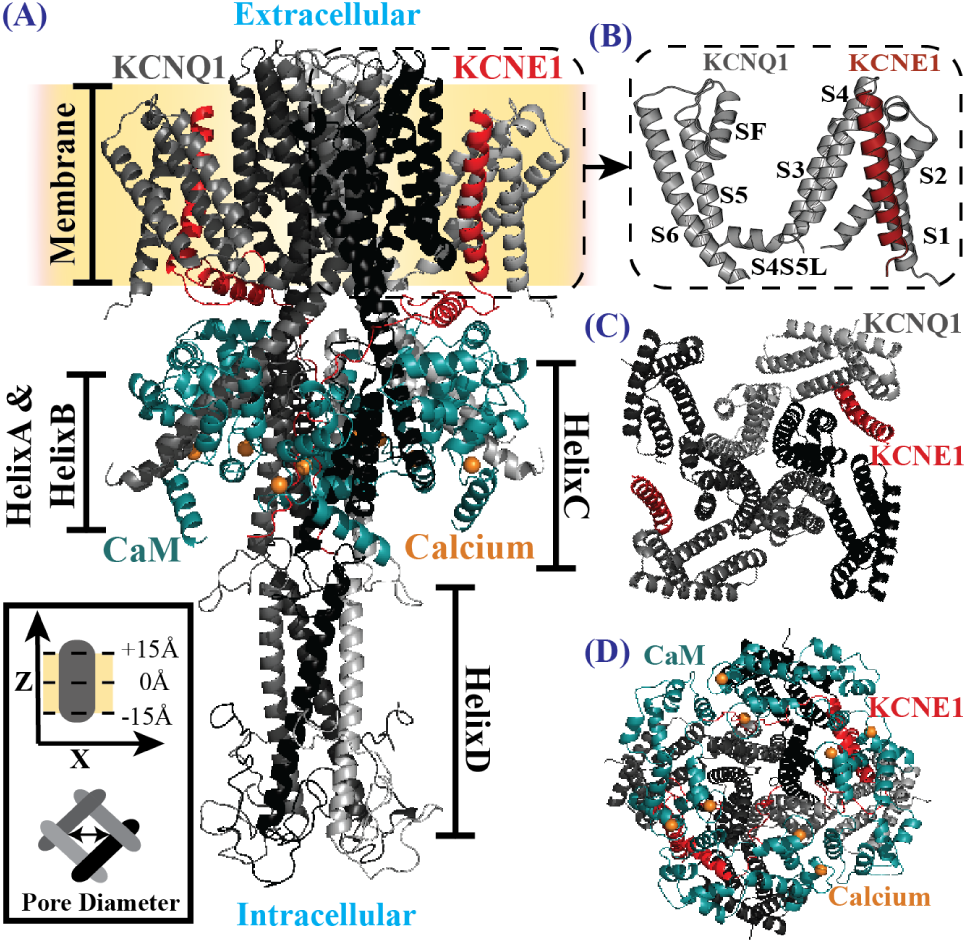
IKs Structure. (A) A side view of initial IKs structure (KCNQ1 tetramer bound to two KCNE1). CaM is bound to HelixA&B and is proximal to the KCNE1 cytoplasmic helix. KCNE1 intracellular segment interacts with HelixC dimer-ofdimers and HelixD tetramer binds with Yotaio (not shown) to facilitate IKs phosphorylation. In addition, HelixC and HelixD are suggested to play a role in subunit dimerization and tetramerization, respectively (26).The inset (bottom-left) shows the coordinate system of IKs (grey; side and bottom views) in the membrane (yellow). (B) An expanded view of the TM S1-S6 arrangement. (C) Top View (without CaM and KCNQ1 C-terminus) and (D) Bottom View of IKs. The structural components and identifying symbols are color coded. *(S4S5L, S4-S5 linker; CaM, Apo-Calmodulin)*

Multidimensional random walks (with Metropolis-Hastings criterion (21)) on the energy landscape at different *V_m_* quantified the transition probability between different regions of the energy landscape. The time-step of the simulated random jumps were estimated from experimentally obtained single-channel kinetic data (9) (~0.17ms). Note that jump resolution could be as small as 1e-12 step and maximum allowed structural deviation between jumps was 1Å. The structures visited in the random walks were clustered in two physiological dimensions of interest – Avg.S4Z (average Z position of 4 VS in the tetramer) and PD (the minimum pore diameter at the activation gate). Conformational jumps between structure pairs from random walks were used to construct the transition matrix of structural clusters (SI Section 2.3). All simulations were conducted at room temperature (21°C) consistent with experiments. Note that PyMOL (22) was utilized for all protein visualizations.

### Calculating SC Using IKs Pore Energy

To simulate IKs function (e.g. single-channel and macroscopic current) starting from simulated structural changes during gating, each structure cluster SC was calculated. A representative structure for each cluster was used to calculate the energy profile (using Particle Mesh Ewald (23)) across the membrane in the pore region (Figure 2A). This pore energy profile (energy barrier) of the representative IKs would affect the probability of an ion passing through the pore (*P_ion_*). The peak of the energy barrier (located at the activation gate) was used to construct the pore energy map (Figure 2A, center). This map was used to obtain parameters, f (an energy barrier scaling constant used to account for extrinsic factors that affect ion conduction) and *G_max_* (maximum conductance), of Eq. 1 and 2, using experimental data (9, 24). A Boltzmann distribution for pore energy (E) was assumed and a self-adaptive differential evolution algorithm (25) was used to estimate SC in Eq. 2. The ionic current (*I_K_*) (Eq. 3) was calculated, based on SC and number of ion channels in cluster i (*N_i_*), with the reversal potential (*E_K_*):

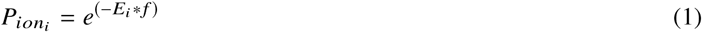

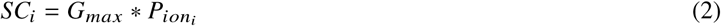

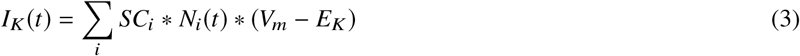

*G_max_* and f were estimated to be 1.9e9 S and 9.31e-2 kT/e, respectively. SC was assumed to be conformation specific and the same at all *V_m_*, because pore energy profile changes due to *V_m_* changes were negligible. Flowchart of simulation steps (SI Fig. S2.1a) and details on specific gating mechanism calculations (SI Section 2.4) are provided in SI.

**Figure 2:**
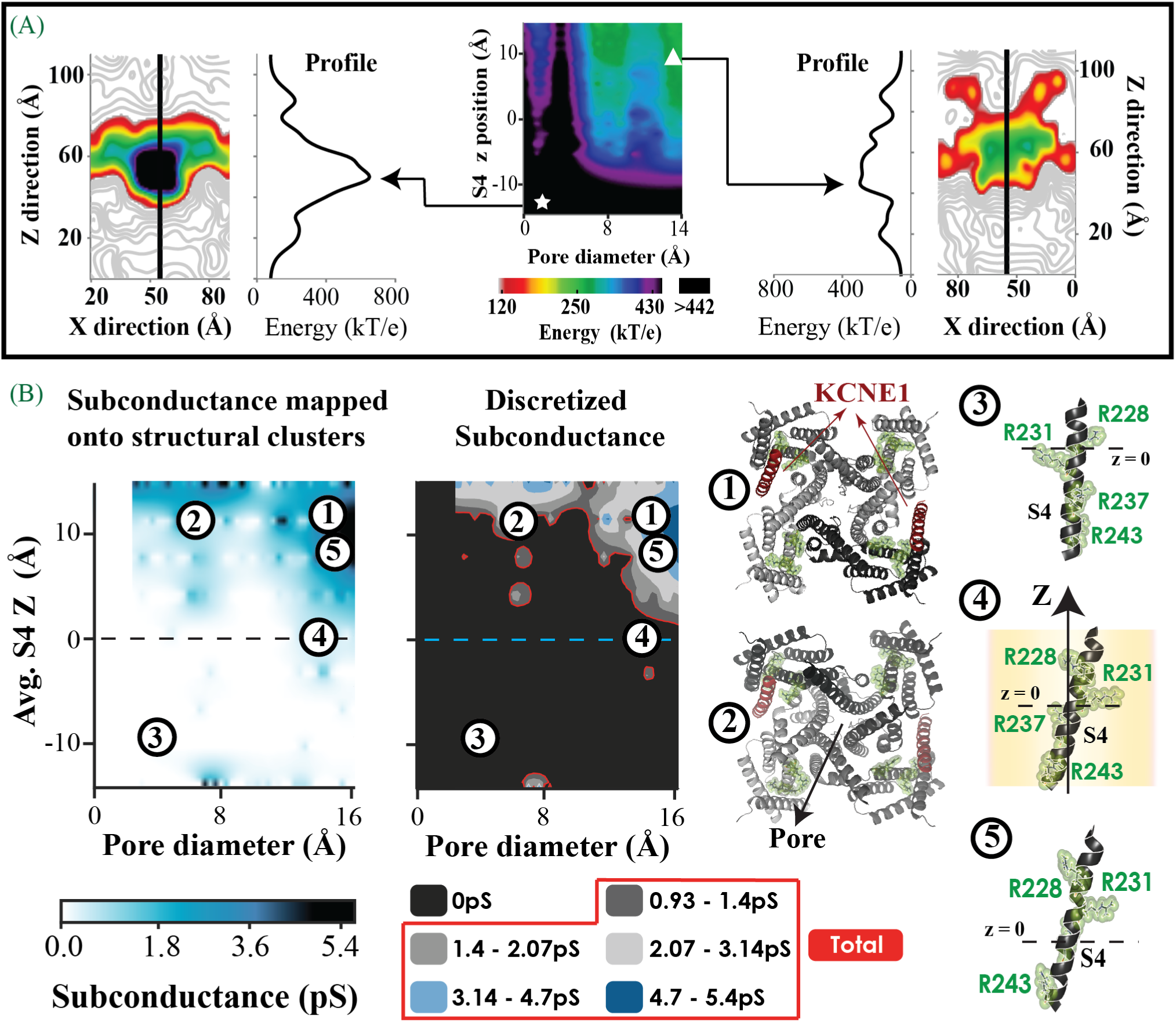
Structure based SC computed from pore energy profile. (A, center) The largest IKs energy barrier across the pore projected on a 2D conformational cluster map (PD and Avg.S4Z position). The symbols indicate representative structures from two different clusters; low S4 Z position and small pore (white star) and high S4 and large pore (white triangle). Corresponding pore energy maps (X-Z slice) for these structures are shown together with the energy profile across the membrane, along a center line (black). (B) The calculated SC values, from pore energy barrier, are projected on a 2D (PD and Avg.S4Z) conformational cluster map (SC map - left). The estimated SC are grouped into five discretized conducting levels, based on experimental measurements (9) (SC map - right). The total conductance (red) is used to simulate macroscopic current and calculate single channel statistics (SI Section 3.1). Protein structures in relation to SC are shown in the right panels. They depict representative pore (bottom view) and S4 (side view) structures from selected clusters on the map. Positively charged S4 residues (R228, R231, R237, R243) are identified (green). Numbers indicate structure locations on the SC map. Only one of four possible S4 conformations is shown.

## RESULTS

IKs structure (Figure 1) was constructed using available experimental data and homology to Kv1.2/2.1 (SI Section 1). Although it was not assumed that S3 and S4 move together, conformations in the library indicate that S3 moves and rotates to accommodate S4 upward movement during gating; this is expected, as the segments are tightly coupled by a short linker. KCNE1 TM segment also moves and rotates to accommodate S4 conformational changes.

### Structural Basis of SC and Simulated Ionic Currents

The pore energy barrier depends on IKs structure. Z-profile of pore energy (Figure 2A) has a large barrier for ions to cross at the pore activation gate. However, this barrier is lower for structures with high Avg.S4Z and large PD (Figure 2A, white triangle). These clusters, with lower energy barriers, have a greater probability of ions passing through the pore as compared to structures with low Avg.S4Z and small PD (Figure 2A, white star). Consequently, they are associated with higher SC levels and greater ionic current. Figure 2B shows the calculated SC map computed from the pore energy map (Figure 2A, center). Structure clusters with the lowest energy barrier (Figure 2A, white triangle) have large PD and high Avg.S4Z; they also have the highest estimated SC (Figure 2B; clusters 1 and 5). As such, SC is dependent on PD as previously suggested (27). Additionally, VS conformational changes and associated IKs structural changes also affect SC.

The estimated SC map (Figure 2B) was used to simulate IKs function as a current carrier. The simulated function was compared to experiments under similar conditions and voltage protocols (9) (Figure 3A-E). Simulated single-channel current traces and the macroscopic current (average of 100 such traces) showed good correspondence with experimental data (Figure 3A,B). SC levels, accessed during activation upon step depolarization to 60mV from −80mV, were equivalent to experimental measurements (Figure 3C). The simulated mean single-channel current amplitudes at different depolarized *V_m_* were used to calculate the microscopic current-voltage (I-V) relationship; an example calculation at 60mV is shown in Figure 3D. The I-V curve shows excellent correlation with experimental recordings (Figure 3E). Similar to experiments, simulations showed many silent single-channel traces for which IKs structural changes did not result in conducting pore conformations. Simulated single-channel latency to first opening during a step depolarization to 60mV was 1.65 ± 0.08s (SI Section 3.1), consistent with the experimental value (1.67± 0.008s, for the current amplitude of 0.5pA). Relevant single-channel current characteristics such as total dwell time, mean open time and 1st opening probability were calculated for each SC level at different *V_m_* (SI Fig. S3.1a,b). Simulated single-channel traces showed increased access to higher SC levels with larger depolarizing *V_m_*.

**Figure 3:**
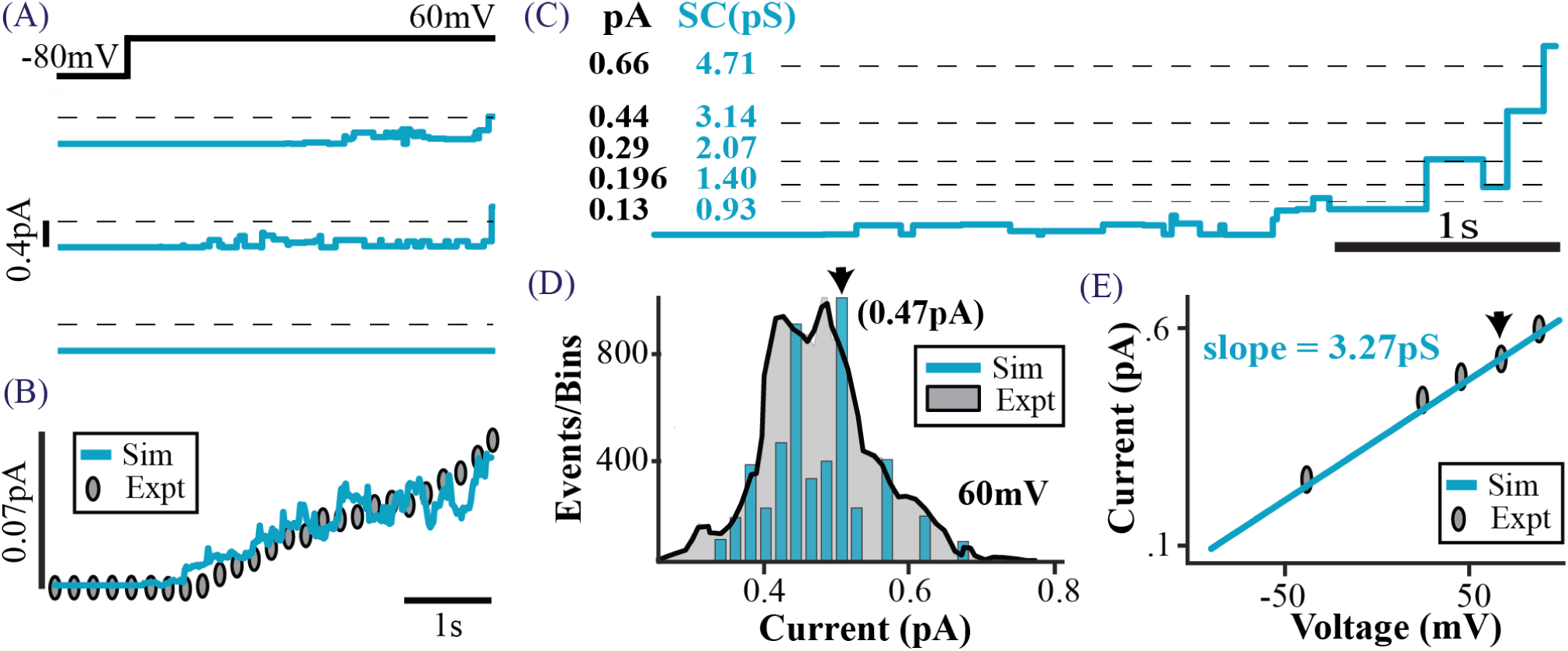
(A) Three simulated single-channel current traces for a step depolarization *V_m_* from −80 to 60mV. (B) Ensemble current (average of 100 traces) fits the experimental data (9). (C) An expanded section of a simulated single-channel trace. Experimentally identified current levels (9) (dashed lines, black entries) and SC computed from simulations (blue entries) are shown. (D) Current amplitude histogram (CAH) of simulated single-channel traces at 60mV (blue) and the corresponding experimental data (shaded grey) (9). The arrow indicates the mean CAH (0.47pA, also indicated by arrow in panel E). (E) Similar histograms (to panel D) are constructed for a range of *V_m_* to obtain the microscopic I-V relationship (blue line) which fits experimental data (symbols) with a slope (mean conductance) of 3.27pS (as determined experimentally (9)).

### Two Distinct VS Movements

Simulated structural changes, associated with VS movement during depolarization (at different *V_m_*), were consistent with results of fluorescence experiments (10). Analysis of 2000 IKs trajectories identified two possible VS movements, at negative and positive *V_m_* (Figures 4). Upon depolarization, VS underwent an initial fast ~9Å upward Z movement followed by a slower ~2Å movement (maximum Avg.S4Z translation at steady-state (SS)). The rate of accompanying PD increase was slow compared to Avg.S4Z translation (Figures 5). Simulated S4 Z movement preceded ionic current, consistent with experiments (10). SS occupation of structural states at −80mV (holding potential, HP) identifies three IKs conformational clusters (HPSS clusters) likely to transition to conducting conformations upon depolarization (Figure 5 (right-top) blue, green, pink symbols). Three likely pathways between HPSS clusters and high SC cluster (Figure 5 (right-top), black symbol) were ascertained for 60mV step depolarization. The temporal probability of each pathway (Figure 5 (right-middle)) showed that conformations with large pore at −80mV transition faster to high SC conformations (large PD, high Avg.S4Z) than conformations with smaller pore. Figure 6 provides a summary and schematic representation of S4 and pore conformational changes during activation.

**Figure 4:**
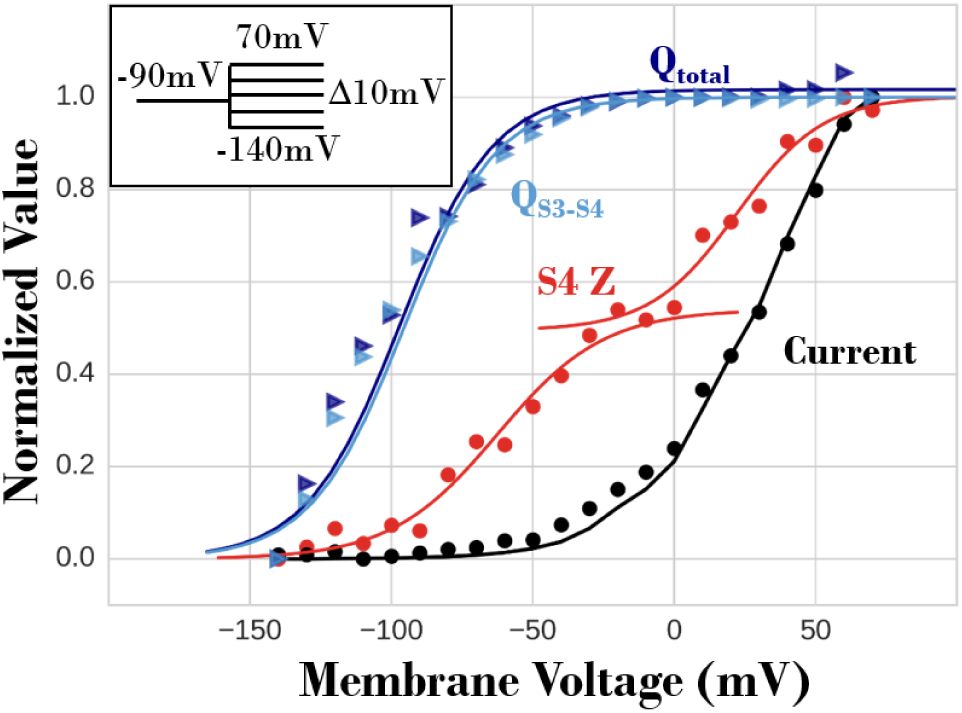
GC displacement (blue), S4 Z (red) and macroscopic current (black). Normalized simulated current (black circles) and sum of tetramer S4 Z displacements (red circles) in 2000 iCh, 4 seconds past depolarization (for protocol in inset). The black line represents experimental data (24). The S4 Z data are fitted with two sigmoidal curves, at negative and positive *V_m_*, representing the I and II movement of S4, respectively. At positive *V_m_*, an increasing fraction of S4 in the 2000 iCh, transition to the II movement. GC displacement associated with one representative iCh S4 Z transitions is calculated based on two selections - the entire IKs (Q_*total*_, dark blue) and S3-S4 segments only (Q_*S*3–*S*4_, light blue) at different *V_m_*. The GC data are fitted with a single sigmoid (light and dark blue traces), analogous to the S4 Z data at negative *V_m_*.

**Figure 5:**
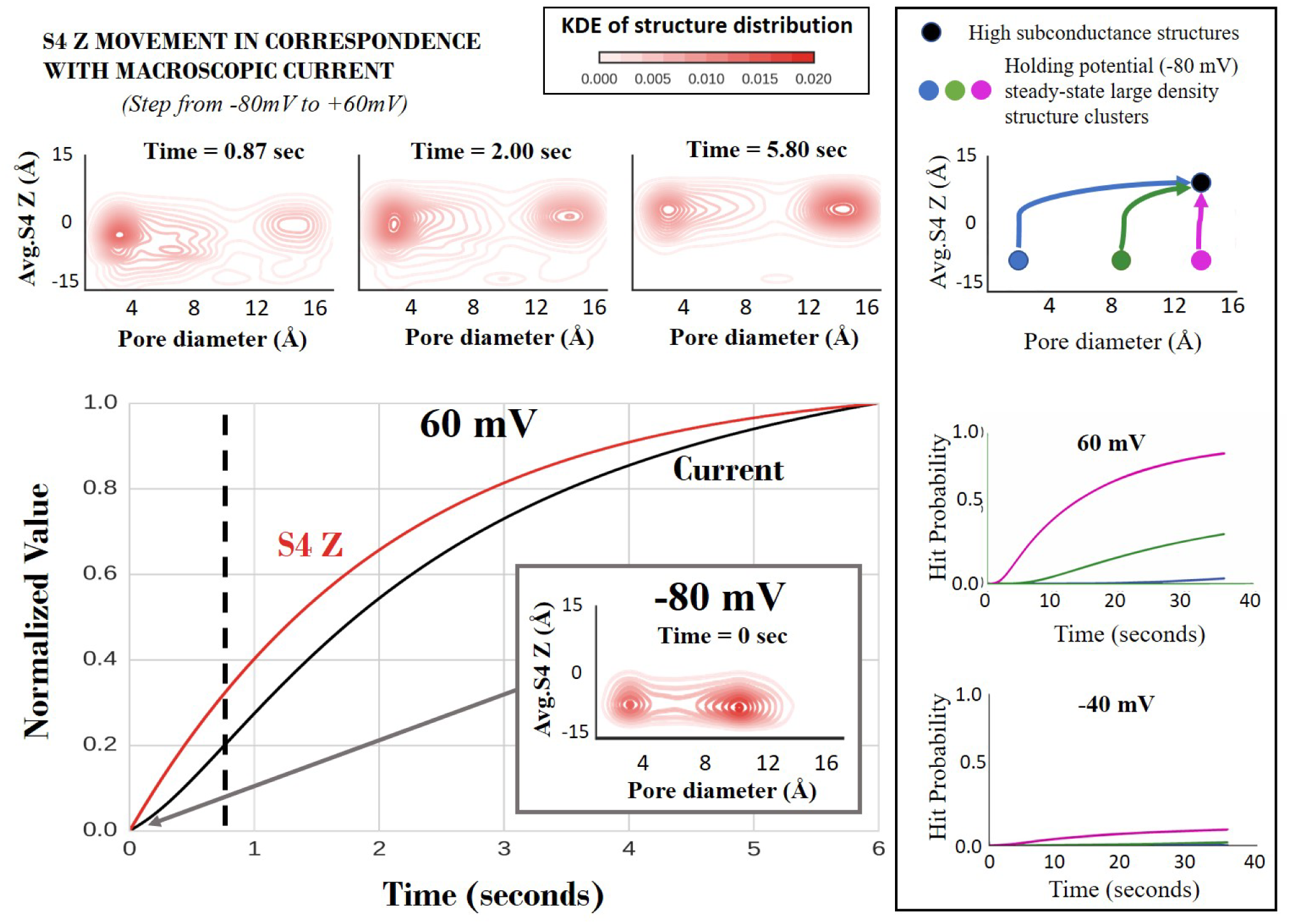
Two movements of the voltage sensor and pore. (Center) The normalized S4 Z movement and macroscopic current (2000 ion-channels), calculated for a depolarizing potential to 60mV from a holding potential of −80mV, shows that the S4 movement in the Z direction precedes ion conduction through the pore. The dashed line divides approximately the two time courses of major conformational changes between structure-clusters shown in the right panel. (Top) The KDE calculated based on the temporal occupancy of the structure-clusters are shown at the top for three different times. Comparing the structure-cluster KDE at −80mV holding potential (center panel (inset)) and 0.87 seconds after 60mV depolarization, it can be seen that the initial movement of S4 in the Z-direction (~9Å) is fast. As time progresses (at 2 and 5.8 seconds), the S4 Z moves slower to a slightly higher Z position (~1.4Å, at 6 seconds) with high conductance. The pore opening from a small to a large pore diameter is also much slower than the initial S4 movement. (Right) The top panel identifies major cluster structures occupied at a holding potential of −80mV (pink, green and blue filled circles). They correspond to the KDE map at −80mV (center panel) except the pink clusters do not show on the map scale. The colored arrows mark the most probable trajectories of the structures to the high SC clusters (black filled circle) during application of a depolarizing potential. The middle and bottom panels show the time progression of the probability of these trajectories (color coded) for two different membrane potentials – 60mV (middle) and −40mV (bottom). *(KDE, kernel density estimate)*

**Figure 6:**
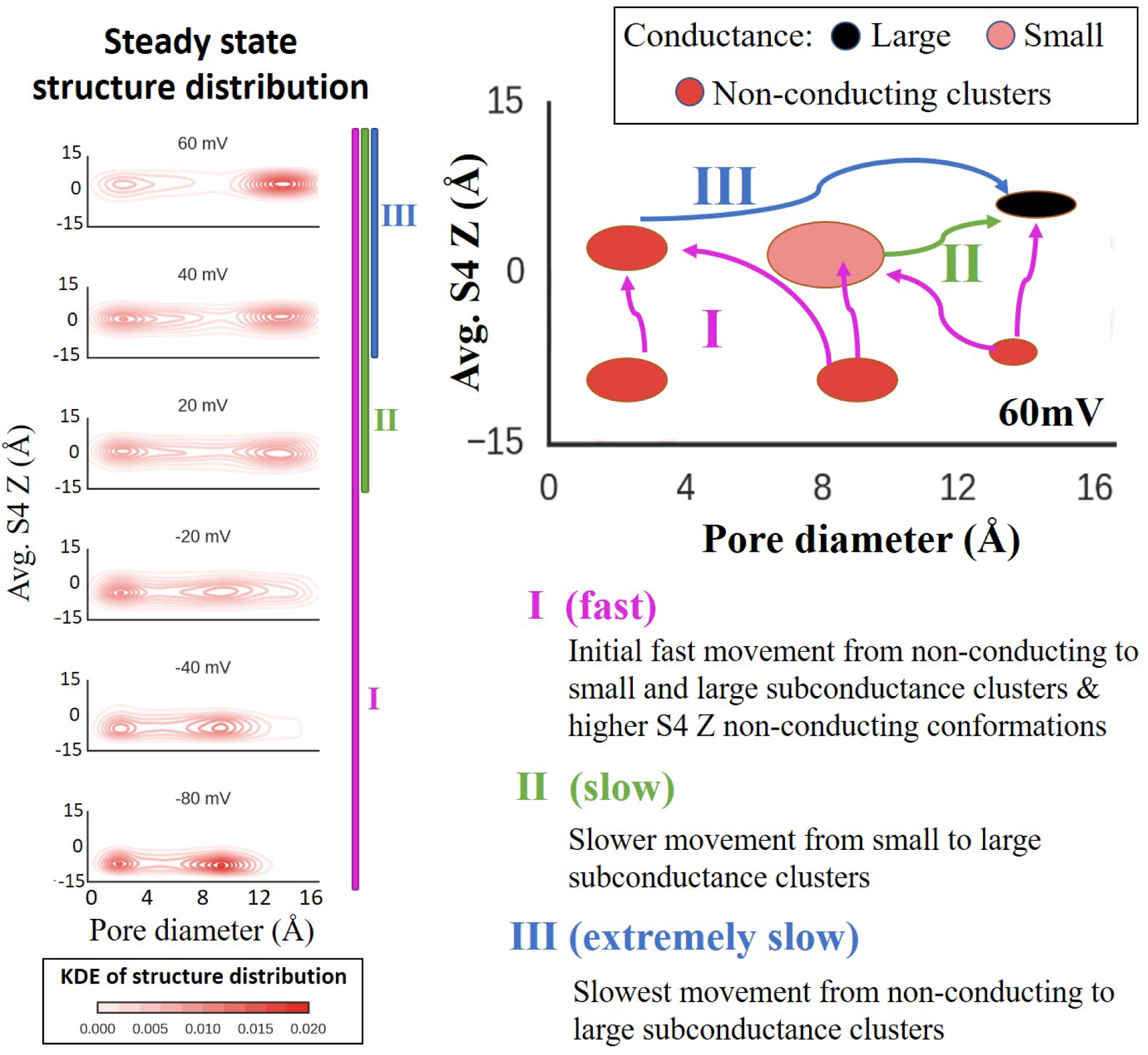
Fast and slow movements of the S4 in Z direction and pore opening. (Left) Steady-state structure cluster KDE for the indicated *V_m_* when depolarized from the holding potential of −80mV. These panels represent transitional structure distributions when a step voltage of 60mV is applied from a holding potential of −80mV. The comparison of the temporal KDE plots (Figure 5, top) with the steady-state KDE (left) indicates that at negative *V_m_* only the first fast transition (I) occurred while at positive *V_m_*, the first transition followed by a slower second transition (II) was observed. The first transition captures the major movement of the S4 in the Z direction, after which the second transition only shows a slight increase in the S4 Z (~1.4Å) and is consistent with the gating charge calculations shown in Figure 4. The slowest transition (III) contributes very little to the current and conformational changes observed in the simulation time course. (Right) Cartoon schema of the fast, slow and extremely slow conformational changes. The pink arrows represent the initial fast conformational transition due to S4 movement in the Z direction, whereas the green arrow shows the slower transition from medium to large pore and the further small S4 Z movement. The blue arrow represents the extremely slow transition from a very small pore to a large pore. Note that the observed transitions are not sequential and happen simultaneously. (KDE, kernel density estimate)

Important residue-pair interactions that affected VS conformational changes were identified, by calculating their contribution to IKs structural energy (SI Fig. S3.2a). There are two S4 positions relative to KCNE1 – proximal and distal (Figure 1C and SI Fig. S3.2a). Proximal S4 charged residues interacted strongly with charged residues at the bottom of the KCNE1 segment (SI Fig. S3.2aC). In contrast, two distal S4 charged residues were only weakly influenced by KCNE1 (SI Fig. S3.2aB). These differences resulted in two types of VS Z movement, of proximal S4 and distal S4, respectively. Residue interaction energy computations (SI Fig. S3.2a) show stabilization of distal S4 at Z position 1.44Å and of proximal S4 at 1.44Å and 2.88Å. In addition, proximal S4 shows strong stabilization at −6.75Å and −4.44Å and increasing stabilization between 0.33 and 2.88Å. These different stabilization profiles (SI Fig. S3.2a-B,C; bottom panel) suggest that distal S4 undergoes a single fast Z movement to its stabilized high Z position, while proximal S4 experiences two movements - a slower Z translation to about −2.78Å followed by a faster upward Z movement to about 3Å. It follows that distal S4 contributes mostly to the initial fast and large Z translation (to ~1Å) and proximal S4 is responsible for the additional slower S4 Z movement (to ~3Å) at positive *V_m_*, resulting in more conformations transitioning to high SC clusters.

### Gating Charge Saturation and Sequential Gating

The model also replicated experimentally observed saturation of GC displacement at positive *V_m_* (Figure 4). This property is the result of the two aforementioned VS Z movements – at negative *V_m_* the VS underwent a fast and large Z translation with large charge displacement, and at positive *V_m_* it experienced an additional slow and small Z translation (Figure 6) that contributed very little to GC displacement. GC calculations with only S3-S4 segments (Q_*S*3–*S*4_) and with the entire protein (Q_*total*_) showed that S3-S4 movement contributes most to GC displacement. Using alternative structural clustering (SI Section 2.4), all possible combinations of S4 Z movements in the tetramer were suppressed and macroscopic current (2000 IKs channels) at the end of 4-second step depolarization was recorded (Figure 7). Ten simulations were conducted for each combination of immobilized S4 and resulting currents were normalized to control (IKs current with all four S4 mobile). If S4 movements were cooperative and required concerted transitions during gating, then immobilization of S4 would have resulted in zero current. However, Figure 7 shows that current decreases linearly with increased number of immobilized S4, consistent with experimental data (11). This linearity confirms independent S4 movement and shows that each S4 contributes incrementally to IKs gating.

**Figure 7:**
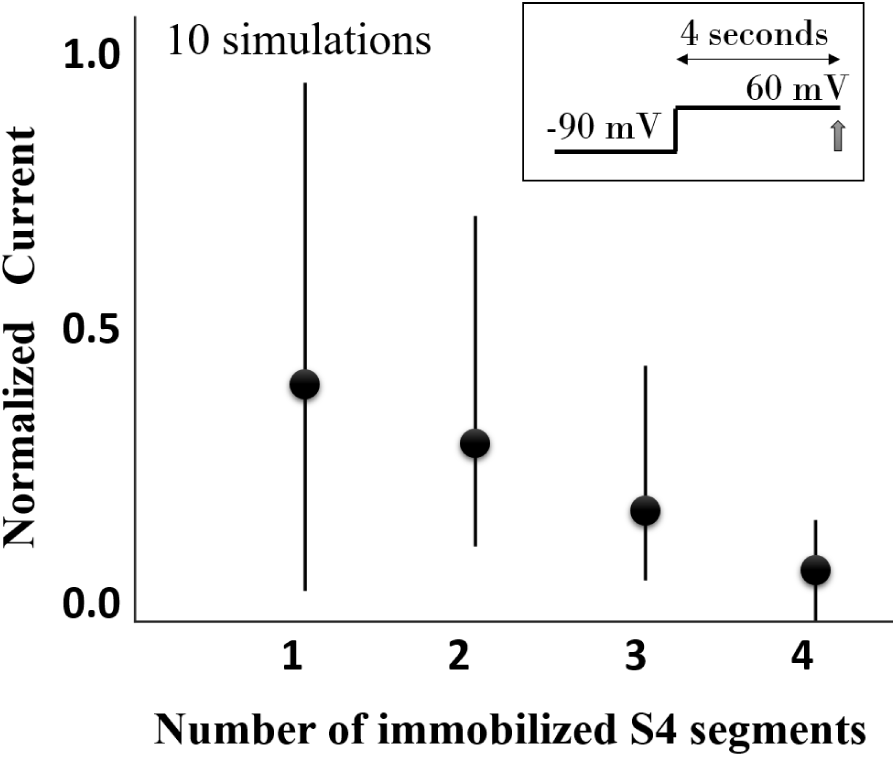
Sequential movement of S4 governs IKs gating. Using an alternative structure clustering method, an increasing number of S4 segments were prevented from moving upward during depolarization to 60mV. The resulting current is measured 4 seconds post-depolarization (protocol in inset). The plot shows a linear decrease of current with increasing number of immobilized S4 segments. Ten simulations were performed for each combination of S4 suppression. For example, two immobilized S4 segments could either be on adjacent or opposite sides of the tetramer. The ten simulations for each of the possible S4 suppression combinations yielded the current mean (filled circle) and the minimum and maximum (error bars) of the plotted values. The results were normalized to the control (simulation of current without S4 movement suppression). The large variance is due to dissimilarity in ionic current reduction depending on whether proximal or distal S4 was immobilized.

### SC Validation

In addition to computing SC from simulated pore energy (Figure 2B), SC were estimated from fitting experimental data (SI Fig. S3.3). The estimated SC map was used to simulate macroscopic current, microscopic current and I-V relationships (SI Fig. S3.3A-D) with high accuracy relative to experiments (9, 24); The SC map from experimental data (SI Fig. S3.3E) was extremely similar to the SC map calculated from pore energy (Figure 2B), with analogous locations of high SC clusters and non-conducting clusters.

## DISCUSSION

Experimental techniques that visualize atomistic structural dynamics (28) are difficult to utilize with large IKs proteins since they contain multiple domains with flexible regions. Molecular dynamics (MD) simulations provide valuable data on a possible structural pathway between two known iCh conformations (endpoints) on the millisecond timescale. Since single-channel dynamics are stochastic, extensive IKs structural changes occur between closed and open states as a function of *V_m_* and time, ensuing in multiple transition pathways over millisecond-to-second timescale. Therefore, multiple endpoints MD computations are not viable on a physiological timescale (5, 29). Previous structure-based functional models did not account for subconductances because a concerted or an allosteric constraint was assumed in order to simulate current (12–15). Although limited by these assumptions, the models could provide a qualitative representation of open probability during KCNQ1 activation. Additionally, the earliest model could also estimate changes to protein energy landscape in the presence of KCNE1 and mutations (15). Knowledge of an experimentally resolved static iCh structure, although insufficient to decipher global structural dynamics, provides a plethora of information (e.g. secondary structure propensities, flexible regions, connectivity etc.) to build a library of possible gating conformations at the required atomistic resolution (Methods and SI - Section 2). Many of our library IKs conformations (sans KCNE1) were found with small deviations from the recently published electron cryo-microscopy (cryo-EM) KCNQ1 structure (SI – Fig. S2.1e).

The ML algorithm (19), used to construct the multidimensional protein energy landscape, converged and was able to predict IKs structural energy with negligible error (SI Section 2.2). This novel approach circumvented the impossible computation of constructing all possible IKs structures. In order to correlate the simulated structural transitions to experimentally measured ionic current, the partition function of the computed energy landscape was obtained from IKs single-channel kinetics (9). Library IKs structural energy shows that electrostatic energy contributes overwhelmingly to the total energy (SI Section 2.1 and Figs S2.1f,g). As a result, changes in IKs energy due to *V_m_* can be adequately represented by changes in electrostatic energy. The simulated structural and functional changes were validated using independent experiments, other than those used for parameter estimations; the IKs model reproduced the recordings of ensemble structural changes, ionic current characteristics and I-V relationships (10, 11, 24).

### IKs structure-based SC determination

IKs structural changes in response to *V_m_* depolarization were simulated and corresponding function (ionic current) was calculated by associating a pore conductance to a structure cluster. Gating conformations with S4 at higher Z have lower energy barriers at the pore activation gate. This is because the repulsive electrostatic field at the pore due to 4 positively charged S4 segments (tetramer) is mitigated by the shielding effect of the surrounding protein segments and water at higher S4 Z positions. Therefore, conformations with low or high S4 Z positions have low or high SC levels, respectively. In addition, SC levels are also affected by PD.

Pore energy profile was computed in the absence of explicit *K*^+^ ions in the pore. This permitted us to calculate SC as an intrinsic property of IKs, independent of extrinsic factors that affect ion conduction such as number, location and hydration characteristics of pore ions. Therefore, SC computations are based solely on IKs molecular structure and its propensity to allow ionic conduction. With these properties, the model correlates structural changes to functional observations and replicates experimentally observed IKs characteristics, including 5 discrete SC levels (between 0.13pS and 0.66pS (9)).

Analysis of early IKs single channel recordings, showed a four times larger conductance as compared to KCNQ1 (24, 30). It also displayed a strong Cole-Moore effect, i.e. activated VS did not result in immediate current. It was suggested that this difference could be due to KCNQ1 ability to access multiple SC states, whereas IKs could only access the largest SC state during activation (31). However, IKs and KCNQ1 do not have access to the same SC states during VS activation. In the absence of KCNE1, the structure-based SC map of KCNQ1 changes significantly. Without the mitigating effect of negatively charged KCNE1 residues, the energy barrier across the KCNQ1 pore is larger for high S4 Z and wide pore KCNQ1 conformations. In the absence of KCNE1, we anticipate that KCNQ1 S4 will behave similar to distal S4 of IKs and will not have the II S4 movement (that allows access to larger SC states; Figure 6). With this information, we envision that both KCNQ1 and IKs conduct ionic current at multiple SC levels and we further predict that the KCNQ1 SC range will be 3-4 times smaller than that of IKs. It follows that, for every addition of KCNE1 to KCNQ1, larger SC states will be accessible by the iCh.

### S4 Movement Precedes Ion Conduction through the Pore

Fluorescence experiments provide data on S4 movement during channel gating. However, this technique cannot quantify the exact S4 movement. Also, depending on the probe utilized, fluorescence intensity measurements can vary. Therefore we assumed that S4 Z translation across the membrane contributes most to changes in fluorescence, with smaller effects due to rotation or X, Y translation. In support of this assumption, the computed sum of four S4 Z displacements (Figure 4) was consistent with experimental fluorescence data (10). As in experiments, simulated macroscopic current showed slower rise relative to S4 Z, indicating that S4 Z movement leads pore ion conduction (Figures 4 and 5). Simulations show that initial movements of S4 and pore, due to depolarizing *V_m_*, result in conformations with low or non-conducting SC. Therefore, little to no current is observed at first. With time, as IKs transitions to higher S4 Z and larger PD, higher SC levels are accessed resulting in a larger current.

Other IKs fluorescence experiments also show two distinct fluorescence–voltage components (F1 and F2); F1 representing the large and fast IKs movements at negative *V_m_* and F2 the smaller and slower component measured at positive *V_m_*, respectively (31, 32). These experiments also showed that the second structural IKs movement (represented by F2) correlated with significant increase in current. Therefore, in agreement with a concerted gating model, it was proposed that F1 and F2 components represented S4 activation and pore opening, respectively (7, 10). However, this IKs gating model could not account for subconductances reported by IKs single-channel recordings and had to be modified to include conductance levels for different numbers of activated S4 in the tetramer (9). Our model is consistent with the latter premise that with increasing number of activated VS, IKs can access larger SC states. However, our model does not reproduce the rapid transitions of IKs from large SC to non-conducting states. We believe that these transitions are caused by extrinsic SC factors that are not represented in this model eg. number and location of ions in the pore, ion hydration, van der Waals interactions between ions, water and pore residues, presence of non-potassium ions in the pore, etc.

### IKs Conformational Trajectories Following Depolarization

Figure 5 (right) identifies important structural clusters and major conformational pathways (groups of trajectories) upon depolarization from −80mV to 60mV and −40mV. The pink pathway encompasses mostly S4 Z translations, and the green and blue pathways include pore expansion. These pathways were identified by analyzing trajectories from each starting HPSS clusters to high SC clusters. Hit probabilities (transition probability between identified clusters) show that very few deeply closed conformations (small pore and low S4 Z) transition to high SC clusters upon depolarization (blue). Interestingly, 78% of total structures occupy this small pore – low S4 Z cluster at −80mV. Very few IKs occupy the pink cluster and thus contribute very little to current upon depolarization. Nevertheless, this trajectory is responsible for the initial current increase. Approximately 18% of conformations occupy the green cluster and contribute to current rise later in time. As expected, the probability of transitioning to high SC clusters is smaller at −40mV than at 60mV. The pathways used to calculate hit probabilities are the major structural transitions that contribute to ionic current during IKs activation. As a model based inference, we suggest that these pathways are closely related to experimentally detected current activation time-constants i.e. the fast and slow time-constants correspond with pink and green pathways in Figure 5, respectively.

### S4 Movement and Pore Opening

Similar to S4 Z upward movement, the pore follows an independent time course of opening. Hit probabilities (Figure 5, right) show that the pathway of mostly S4 upward movement (pink) was much more probable than other pathways (green and blue). The pathway that required the largest PD increase (blue) to reach high SC states was least probable. Thus, initial Avg.S4Z transition (from −10 to ~1Å) was much faster than the second Avg.S4Z transition (from Ȉ1 to 3Å), which in turn was faster than pore opening. A cartoon in Figure 6 (right) explains these dynamics further.

Using KDE (kernel density estimation) of occupied structure clusters at SS, for a range of *V_m_* between −80 and 60mV, helped identify important cluster groups that were visited upon depolarization (Figure 6, left). Temporal KDE data (Figure 5, top), show that conformations experiencing step depolarization to negative *V_m_* undergo only the first fast transition (I) where the dynamics are mostly determined by upward S4 Z translation (from −8Å to ~1Å). For positive depolarization to *V_m_*, conformations initially experience the first transition followed by a second slower transition (II) of small S4 Z movement (from ~1Å to 3Å) and medium to large PD increase, consistent with experimental data (10). The VS two-step transitions correlate with the fast (I) and slow (II) transitions of S4 (Figure 4). Ionic current follows S4 movement with a delay, because pore opening is slow compared to S4 Z transitions.

The third much slower transition (III), from very small to large pore, involves small S4 Z movement (from ~1Å to 3Å) that occurs at positive *V_m_* and takes much longer time. Consequently, IKs channels take very long time to reach SS. This provides a likely explanation for the experimentally observed IKs current increase over very long time post-depolarization.

For one iCh, I contributes most to GC displacement. II, at positive *V_m_*, involves a very small S4 Z translation and therefore contributes much less to GC displacement. As *V_m_* increases, more iCh transition to conformations with higher S4 Z, as reflected in the sum shown in Figure 4 (red). However, for one iCh, the S4 Z movement is small and contributes minimally to GC, which exhibits saturation (Figure 4, blue).

Our structurally-based model of IKs shows similar characteristics to the Zaydman-Cui kinetic (Markov) IKs model (32); the movement ‘I’ (resting to intermediate state transitions) confers little to no current in both models. It follows that our structure-based model can be correlated to the Zaydman-Cui kinetic model that represents simplified transitions from resting to intermediate and intermediate to activated states. The correspondence requires a small modification to the Zaydman–Cui model of IKs; instead of suppressing transitions from intermediate-closed to intermediate-open states in the presence of KCNE1, the intermediate-open and activated-open states could be associated with very small and large conductances, respectively. This adjustment will enable transitions between intermediate-open and activated-open states, vital for IKs activation in our structure-based model. With this modification, the resting, intermediate and activated kinetic states can be associated with Avg.S4 Z conformations less than −5Å, between −5Åand ~1.2Åand greater than ~1.2Å, respectively. Also, we suggest that the Zaydman-Cui model of KCNQ1 could be modified with similar adjustments, but with a reduced difference between intermediate-open and activated-open conductances, to account for its smaller SC range. The Zaydman-Cui model suggests that the functional VS-pore coupling is reduced in IKs relative to KCNQ1. Our structure-based model of IKs supports this supposition, as transitions to intermediate conformations result in extremely small current, and transitions to activated states are required for substantial increase in current.

### Residue Interactions that Govern S4 Z Movement

IKs conformational changes, in the presence of *V_m_*, depend largely on the environment of charged S4 residues. SI Fig. S3.2aA shows a cartoon depicting S4 charged residues, as well as the surrounding charged residues in the voltage sensor domain (S1-S4) and KCNE1 TM segment. Interaction strength between residue pairs and contribution of individual charged S4 residues to protein stabilization show that proximal and distal S4 undergo two types of movement, based on their proximity to KCNE1. Distal S4 is likely to move upward faster than proximal S4 upon depolarization (SI Fig. S3.2aB) because strong interactions between proximal S4 and KCNE1 slow S4 transitions from a low to high Z positions, resulting in slow activation (SI Fig. S3.2aC). Charged KCNE1 residues (R67, K69, K70 and E72) interact strongly with proximal S4 residues and have weaker interactions with distal S4 residues. These KCNE1 residues contribute most to proximal S4 stabilization at low Z through their interactions with R243 and D242 residues. Distal S4 is likely to be stabilized at ~1Å after depolarization and is unlikely to transition further in the Z-direction. However, proximal S4 (although slower), can transition to higher S4 Z positions (~3Å). Ensemble VS movement has an initial large transition (fast) followed by a smaller Z transition (slow). The second ensemble S4 movement, observed at positive *V_m_* (Figure 4), is likely the result of proximal S4 transitioning from intermediate to high Z positions. The first ensemble S4 movement is possibly a combination of fast distal S4 (low to high) and slow proximal S4 (low to intermediate) Z transitions. In both proximal and distal S4, R228 plays an important role in stabilizing S4 at high Z positions.

### Sequential Gating

The simulations show that macroscopic current increases linearly with increasing number of moving S4 segments (Figure 7). This is because IKs SC depends on all four S4 and pore conformations. High SC conformations are only accessible when all four S4 move to high Z positions with large PD. Nonetheless, small and intermediate SC conformations are accessible at low S4 Z with medium to large PD. IKs pore is not tightly coupled to S4 Z movement because of the flexible S4-S4S5L three-residue segment (QGG; an additional ~25° flexibility compared to Kv1.2/2.1 (33)). Thus, IKs pore openings (with small or intermediate SC conformations) are possible at lower S4 Z conformations. Large pore opening is only possible with all four S4 at high Z. Structurally, the pore can open (increase in diameter) at low-to-medium S4 Z positions, but the model IKs does not present with significant current until S4 Z reaches relatively high positions. Consequently, from a model based inference, we suggest that IKs gating is structurally allosteric but functionally concerted.

## CONCLUSIONS AND LIMITATIONS

The methodology presented here is a computational framework for studying, through simulations, atomistic interactions and dynamics of a protein that govern its biological function. The framework is applicable to most proteins and is a powerful tool for understanding protein behavior. This methodology can also be applied to study the effects of mutations, ligands and drugs. Additionally, IKs structural and functional changes during an AP can be examined (34, 35).

The structural model (Figure 1) is not based on the recently published cryo-EM data of KCNQ1 structure bound to calmodulin (CaM) (33). This cryo-EM structure presents close homology to Kv1.2/2.1 TM segments and is very similar to the initial IKs structure in our library, with some differences in the intracellular domain. In particular, our model presents a different S6-HelixA attachment and number (3 in Cryo-EM vs. 2 in model) of Ca ions bound to CaM. The framework presented here allows incorporating these features into future structural libraries for evaluation of their effects on gating. We estimate that contribution to IKs energy will be minimal, as CaM is in the water/ion medium. Also, the presence of KCNE1 in our structures (but not in the cryo-EM structure) is an important feature of human IKs as it could restrict pore opening.

The presented model uses implicit dielectrics to represent membrane and water/ion surfaces. Thus, it is possible that important van der Waals interactions between the protein and membrane, that affect gating, could have been underestimated. However, given that IKs C-terminus extends over 60Å, explicit calculations would slow the computation 200-fold. With future advances in computational hardware and algorithms, it will likely be possible to incorporate an all-atom energy calculation as part of the gating model. In this study, protein dynamics were coarse-grained (by clustering structures) but was sufficiently fine to replicate all experimental data. The resolution can be tuned, based on study requirements, by increasing the number of structure clusters and random walks.

## AUTHOR CONTRIBUTIONS

S.R. performed the simulations and analyzed data; Y.R. supervised the project and analyzed data. Both authors conceived the project and wrote the manuscript.

## ACKNOWLEDGMENTS

The authors are grateful to Prof. Bernard Attali, Tel Aviv University, for providing experimental data. This study was supported by NIH–National Heart, Lung, and Blood Institute grants R01-HL-033343 and R01-HL-049054 (to Y. Rudy). Dr Rudy is the Fred Saigh Distinguished Professor at Washington University. Computations were performed using facilities in the Rudy Lab and Washington University Center for High-Performance Computing, which is partially supported by grant NCRR 1S10RR022984-01A1.

## SUPPLEMENTARY MATERIAL

An online supplement to this article can be found by visiting BJ Online at http://www.biophysj.org.

